# Unbiased profiling of CRISPR RNA-guided transposition products by long-read sequencing

**DOI:** 10.1101/2021.02.11.430876

**Authors:** Phuc Leo H. Vo, Christopher Acree, Melissa L. Smith, Samuel H. Sternberg

**Author notes:** Corresponding author: Samuel H. Sternberg. **Email:**.

## Abstract

Bacterial transposons propagate through either non-replicative (cut-and-paste) or replicative (copy-and-paste) pathways, depending on how the mobile element is excised from its donor source. In the well-characterized *E. coli* transposon Tn7, a heteromeric TnsA-TnsB transposase directs cut-and-paste transposition by cleaving both strands at each transposon end during the excision step. Whether a similar pathway is involved for RNA-guided transposons, in which CRISPR-Cas systems confer DNA target specificity, has not been determined. Here, we apply long-read, population-based whole-genome sequencing (WGS) to unambiguously resolve transposition products for two evolutionarily distinct transposon types that employ either Cascade or Cas12k for RNA-guided DNA integration. Our results show that RNA-guided transposon systems lacking functional TnsA primarily undergo copy-and-paste transposition, generating cointegrate products that comprise duplicated transposon copies and insertion of the vector backbone. Finally, we report natural and engineered transposon variants encoding a TnsAB fusion protein, revealing a novel strategy for achieving RNA-guided transposition with fewer molecular components.

---

DNA transposons are ubiquitous mobile genetic elements (MGEs) found in all domains of life that encode transposase enzymes necessary for their spread within and between genomes (1, 2). Until recently, transposases were thought to catalyze either highly promiscuous transposition, generating random insertions with little sequence specificity at the target site, or highly selective transposition to conserved genomic sequences (termed “attachment sites”), or transposition with variable degrees of target specificity, including preferences for short degenerate motifs or bent DNA; in all cases, the targeting pathways are dependent exclusively on protein-DNA interactions (3). This paradigm was upended with the discovery that multiple types of bacterial transposons employ nuclease-deficient CRISPR-Cas systems for programmable transposition, in which target sites are defined by RNA-guided DNA binding (4–6). In the case of Tn*7*-like CRISPR-transposons (CRISPR-Tn), a multisubunit protein-RNA complex encoded by type I CRISPR-Cas systems, called Cascade, directs high-fidelity DNA integration (**Fig. 1a**) (5, 7–9). In the case of Tn*5053*-like CRISPR-transposons, DNA integration is guided by RNA and Cas12k, a single-effector protein encoded by type V-K systems, though genome-wide specificity is much lower (6, 8, 10). These findings highlighted the molecular versatility of RNA-guided DNA targeting, and reveal how CRISPR-Cas functionality has been repurposed for MGE transmission.

**Figure 1.**
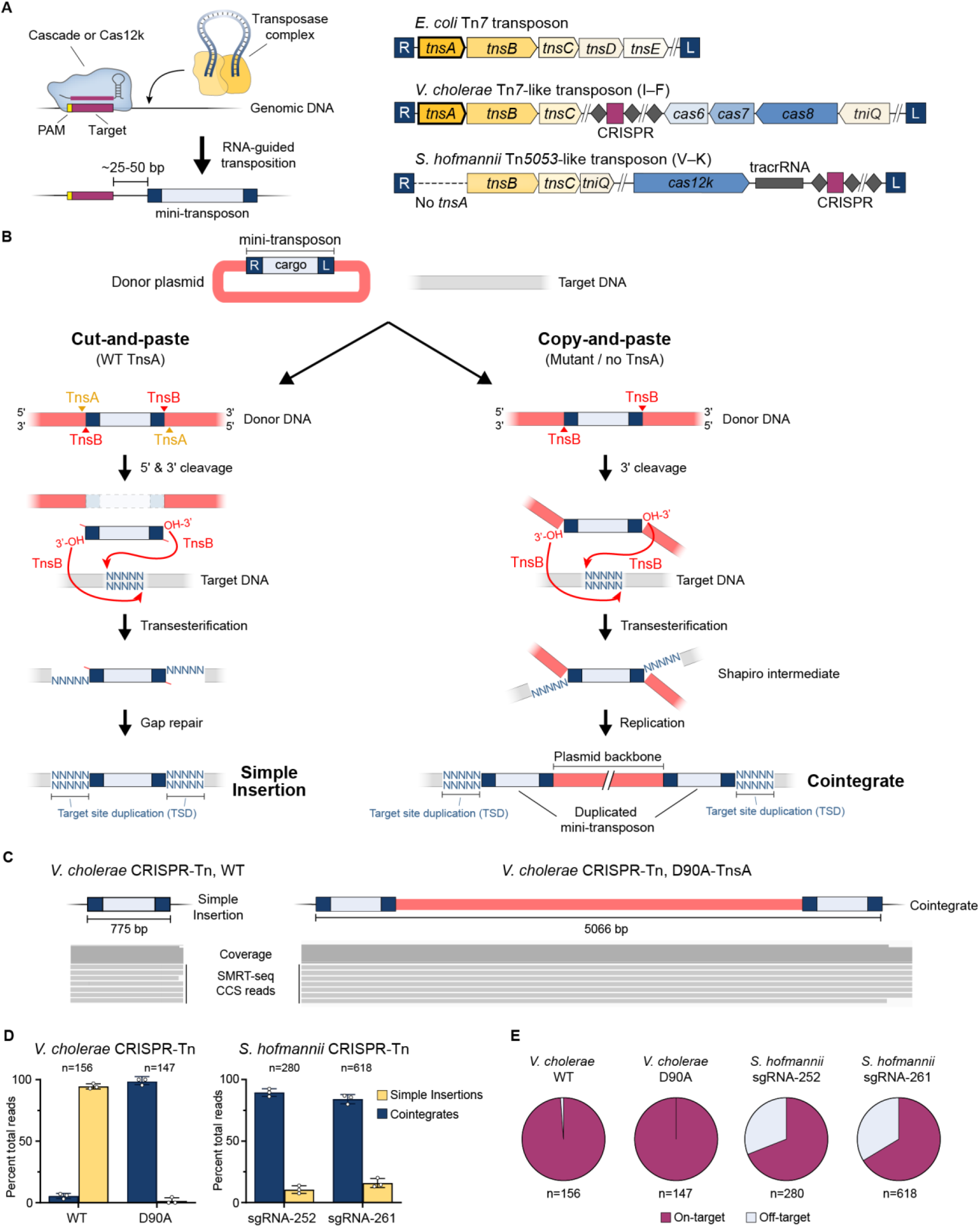
Whole-genome SMRT-sequencing resolves simple insertion and cointegrate transposition products. **a,** Left, general mechanism of RNA-guided transposition directed by Cascade (Type I) or Cas12k (Type V). Right, genetic architecture of the *E. coli* Tn*7* transposon, and Tn*7*-like or Tn*5053*-like transposons mobilized by type I-F or type V-K CRISPR-Cas systems, respectively. Note that Tn*5053*-like transposons do not encode TnsA. **b,** Roles of TnsB and, when present, TnsA, in DNA excision and integration. Non-replicative cut-and-paste transposition involves both 5’ (TnsA) and 3’ (TnsB) cleavage of the donor DNA, resulting in simple insertion of the mini-transposon (left). Replicative copy-and-paste transposition involves only 3’ cleavage, resulting in a Shapiro intermediate and eventual cointegrate product containing duplicated mini-transposon and embedded plasmid backbone (right). **c,** Representative SMRT-seq CCS reads from the wild-type *V. cholerae* CRISPR-Tn show hallmarks of simple insertion products (left); SMRT-seq reads from a D90A-TnsA mutant show hallmarks of cointegrates (right). **d,** Population-level quantification of simple insertion and cointegrate transposition products from *V. cholerae* CRISPR-Tn using either WT TnsA or D90A-TnsA mutant (left), and the WT *S. hofmannii* CRISPR-Tn programmed by two distinct guide RNAs (right). Data represent mean ± s.d. for 3 biological replicates; n denotes the total number of CCS reads. **e,** Specificity of RNA-guided transposition determined from SMRT-seq data, for both *V. cholerae* and *S. hofmannii* CRISPR-Tn systems. Data are shown as mean for 3 biological replicates; n denotes the total number of CCS reads.

While there is extensive literature describing the biochemical mechanisms of transposition for well-studied bacterial transposons such as Mu, Tn*3*, and Tn*5* (11), our understanding of CRISPR-Tn mechanisms is largely extrapolated from prior work on the homologous, non-CRISPR transposon from *E. coli*, Tn*7* (12). As Tn*7* employs TnsA and TnsB to fully excise the transposon DNA via cleavage of both strands at both transposon ends, thereby facilitating non-replicative (cut-and-paste) transposition (13, 14), we previously hypothesized that Tn*7*-like CRISPR-Tn systems would follow the same pathway (5) (**Fig. 1a**). In contrast, Tn*5053* encodes only TnsB, and was previously shown to direct replicative, copy-and-paste transposition (15). Based on these data, it could be reasonably anticipated that type V CRISPR-Tn systems encoding homologous transposase proteins should similarly undergo copy-and-paste transposition, resulting in cointegrate products with insertion of the entire donor DNA molecule and a duplicated transposon (**Fig. 1b**). However, a previous study of a type V CRISPR-Tn system (ShCAST), relying solely on junction-PCR approaches, failed to detect the existence of cointegrates and proposed an incomplete transposition model (6, 16, 17).

To unambiguously resolve the full-length products of RNA-guided transposition, we adopted PCR-free, single-molecule real-time sequencing (SMRT-seq) and characterized multiple, evolutionarily distinct transposon types. Our results demonstrate that the presence or absence of functional TnsA dictates whether transposition occurs via non-replicative or replicative pathways, respectively, and further show that cointegrate products generated by a Tn*5053*-like CRISPR-Tn are frequently off-target and resolve inefficiently via recombination. Our study highlights the importance of carefully considering the wealth of transposition biochemistry when exploring new MGEs, and reveals functional consequences of transposons that harbor fewer molecular components. We also report the first example of an active CRISPR-Tn that encodes a single, multifunctional TnsAB fusion transposase, and show that this minimal system generates high-fidelity, RNA-guided DNA insertions.

## RESULTS

We previously discovered and optimized a *Vibrio cholerae* CRISPR-Tn (Tn*6677*) for RNA-guided DNA integration in a heterologous *E. coli* host (5). Using a two-plasmid expression system (**Methods**), we transformed *E. coli* cells and targeted the genome for integration using crRNA-13. Unlike our previous SMRT-seq experiments, in which individual clones were analyzed (8), here we pooled a large population of transformants, isolated gDNA, performed WGS, and carried out downstream analyses using circular consensus sequence (CCS) reads (**Supplementary Fig. 1**). Given the >11 kb average CCS read lengths (**Supplementary Table 1**) and unbiased library preparation, we reasoned that transposon-containing reads would unambiguously report the type and genetic structure of integration products sampled across the entire population.

The majority of transposon reads with the wild-type system corresponded to on-target integration events (~50-bp downstream of the target site) and bore hallmark features of non-replicative transposition, in which the simple insertion was flanked by genomic sequences and the expected 5-bp target site duplication (TSD) (**Fig. 1c**). In contrast, when we analyzed gDNA from cells expressing a TnsA D90A mutation, which is predicted to abrogate 5’-strand cleavage (5, 13), the raw reads instead bore hallmark features of replicative transposition, in which the entire pDonor vector backbone was integrated in the genome, flanked by duplicated copies of the mini-Tn (**Fig. 1c**). By selectively analyzing only those reads spanning both transposon ends to unambiguously classify the read as either deriving from pDonor, a simple insertion, or a cointegrate (**Methods**), we next quantified the ratio of simple insertion versus cointegrate products across the population. Whereas the WT system predominantly generated simple insertions, over 95% of reads from the TnsA mutant system were consistent with cointegrate products (**Fig. 1d**).

We next investigated Tn*5053*-like CRISPR-transposons, which notably lack *tnsA* and are guided by the Cas12k-sgRNA module (6). Upon testing two different sgRNAs for a WT *S. hofmannii* CRISPR-Tn described previously (6), we found that >85% of transposon-containing reads were consistent with cointegrate products (**Fig. 1d**). Interestingly, whereas >98% of SMRT-seq reads for the *V. cholerae* CRISPR-Tn represented on-target DNA integration, only 65-70% of SMRT-seq reads for the *S. hofmannii* CRISPR-Tn were on-target (**Fig. 1e**, **Supplementary Fig. 2, Methods**), consistent with our prior analysis of genome-wide integration specificity (8). These results conclusively demonstrate that CRISPR RNA-guided transposons lacking functional TnsA are unable to fully excise from their donor sites as a linear species, and instead mobilize through a copy-and-paste pathway that involves replication through the transposon and insertion of the entire donor plasmid. Although many transposon cointegrates resolve through the action of dedicated recombinase proteins, also known as resolvases (18, 19), Tn*5053*-like CRISPR-transposons do not encode an identifiable recombinase. Thus, simple insertions would be expected to arise only through a homologous recombination-dependent pathway requiring the combined action of dedicated host DNA repair factors.

From a technology perspective, Cas12k-directed transposons offer ostensible advantages for RNA-guided DNA integration because of their reliance on fewer molecular components and smaller coding size, albeit with considerable drawbacks in product purity. In the course of our efforts to engineer more streamlined RNA-guided transposons that maintain high-fidelity activity, we uncovered a CRISPR-Tn subtype that harbors a novel gene encoding a fusion TnsAB polypeptide, with clearly identifiable endonuclease- and DDE integrase-family domains (**Fig. 2a**). After selecting a candidate transposon from *Aliivibrio wodanis* for gene synthesis, we found that this system was fully capable of RNA-guided DNA integration, and exhibited similar properties as the *V. cholerae* CRISPR-Tn (**Fig. 2b**). Next, we designed synthetic fusions of *V. cholerae* TnsAB modeled off of the *A. wodanis* protein, and showed that these variant systems have wild-type integration activity (**Fig. 2c**). When we profiled transposition products from these fusion systems by SMRT-seq, we found that the WT and TnsA-mutant systems generated predominantly simple insertions or cointegrates, respectively, with the WT TnsAB fusions producing almost no detectable cointegrates across a population of cells (**Fig. 2d,e**). Finally, on-target analysis of SMRT-seq reads showed that these systems maintained the high target specificity seen in WT *V. cholerae* CRISPR-Tn (**Fig. 2f**, **Supplementary Fig. 2**). These findings reveal a promising avenue for reducing the number of biomolecules necessary to achieve RNA-guided DNA integration in heterologous organisms.

**Figure 2.**
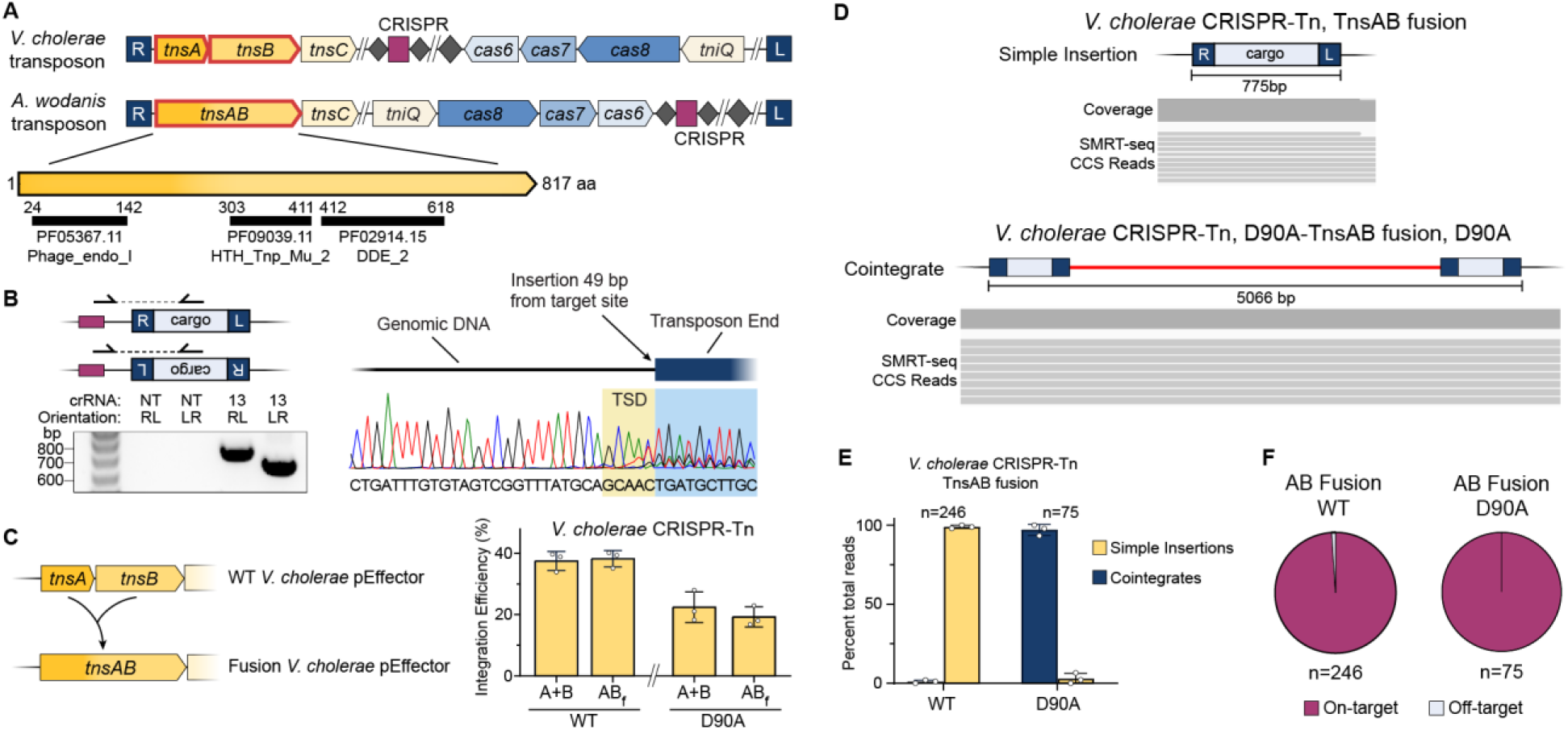
RNA-guided transposition with Tn*7*-like CRISPR-transposons encoding TnsAB fusion proteins. **a,** Genetic architecture of *A. wodanis* transposon harboring a natural *tnsAB* fusion gene, with pfam domains for the TnsAB protein shown below. **b,** PCR analysis and Sanger sequencing confirm RNA-guided DNA integration with *A. wodanis* CRISPR-Tn. **c,** qPCR analysis showing integration activity with the *V. cholerae* CRISPR-Tn, for pEffector encoding either TnsA and TnsB (A+B) or a fusion TnsAB polypeptide (AB_f_), with WT or D90A mutation. Data are shown as mean ± s.d. for 3 biological replicates. **d,** Representative SMRT-seq reads from the *V. cholerae* CRISPR-Tn with fused TnsAB, showing hallmarks of simple insertion products (top); SMRT-seq reads from the same system with D90A-TnsAB are shown at bottom, showing hallmarks of cointegrate products. **e,** Population-level quantification of simple insertion and cointegrate transposition products from the *V. cholerae* CRISPR-Tn with TnsAB fusion, for either the WT system or a D90A-TnsAB mutant. Data represent mean ± s.d. for 3 biological replicates; n denotes the total number of CCS reads. **f,** Specificity of RNA-guided transposition determined from SMRT-seq data, for the *V. cholerae* CRISPR-Tn with TnsAB fusion. n denotes the total number of CCS reads.

## Discussion

Determining the genome-wide effects of DNA editing reagents in a rigorous and unbiased fashion is of paramount importance for safe and effective deployment of CRISPR-based technologies (20). Indeed, the observation biases imposed by previous analyses of CRISPR-Cas9 and CRISPR-transposons have obscured critical, unanticipated byproducts such as large deletions or vector insertions (6, 16, 21, 22). By applying SMRT-WGS, in conjunction with prior transposon-insertion sequencing approaches (5, 8), we have now exhaustively characterized the types and sites of genome-wide transposition products afforded by Tn*7-* and Tn*5053*-like CRISPR-transposons.

The Tn*7*-like *V. cholerae* CRISPR-Tn produces primarily simple insertions, whereas mutation of the TnsA active site results in cointegrates. Interestingly, we still observe rare cointegrate formation with WT TnsA, which may result from asynchronous donor cleavage by TnsA and TnsB, cut-and-paste transposition from low-abundance dimeric donor plasmids, or homologous recombination between simple insertion products and the cellular reservoir of free donor plasmid. When TnsA is completely absent, as in the Tn*5053*-like *S. hofmannii* CRISPR-Tn system, cointegrates are the major products of transposition. Nevertheless, we still observed low-frequency simple insertion products, which may result from events in which the 5’ transposon ends are nicked by host enzymes, or from homologous recombination between duplicated transposon copies.

Overall, our data confirm the decisive role of TnsA in mediating non-replicative, cut-and-paste transposition across diverse systems. This key function, alongside the ability of TnsA and TnsB to function as a fusion polypeptide in Tn*7*-like CRISPR-Tn systems, hint at the future potential to engineer TnsAB fusions into Tn*5053*-like CRISPR-Tn systems as a synthetic strategy to produce high-purity simple insertion products. The TnsAB fusion we describe highlights new strategies for streamlining delivery and/or expression of the multiple polypeptide components necessary for applications of RNA-guided DNA integration technology in heterologous cell types.

## METHODS

### Constructs and transposition assays

Transposition assays were performed by co-transforming *E. coli* with two plasmids: an effector plasmid (pEffector) encoding the guide RNA and all protein components on a pCDFDuet-1 backbone, and a donor plasmid (pDonor) encoding a ~1-kb mini-transposon on a pBBR1 backbone. pEffector plasmids for the *V. cholerae* (Vch INTEGRATE - Tn*6677*) and *S. hofmannii* (ShCAST) systems were cloned from versions described previously (8). TnsA-D90A pEffector plasmids for the *V. cholerae* system were generated by introducing a D90A mutation in TnsA. TnsAB fusion pEffector plasmids for the *V. cholerae* system were cloned by inserting a 5’-GC-3’ sequence in the region where the TnsA and TnsB open reading frames overlap. The D90A version of the TnsAB fusion protein was generated by introducing the D90A mutation in the TnsAB fusion protein. Components for the *A. wodanis* system were derived from *Aliivibrio wodanis* strain 06/09/160 genome (NCBI accession LR721750.1), synthesized as fragments (GenScript), and cloned into the pCDFDuet-1 backbone under control of an IPTG-inducible T7 promoter.

Transposition assays for the *V. cholerae* and *S. hofmannii* systems were performed by transforming chemically competent WT BW25113 *E. coli*. Cells were recovered at 37 °C for 1 hr in LB media, plated on LB-agar with appropriate antibiotic selection, and incubated for 30 hr at 37 °C. Transposition assays for the *A. wodanis* system were performed similarly by transforming BL21(DE3) *E. coli*, with final incubation on LB-agar also containing 0.1 mM IPTG for expression induction.

### PCR and qPCR analyses

Colonies were scraped after incubation, resuspended, and subjected to heat lysis. PCR/qPCR analysis of the resulting lysates were performed as described previously (5). Briefly, cells were resuspended in water, lysed at 95 °C for 10 min, and pelleted at 4000*g* for 2 min. For each sample, the resulting supernatant was diluted 20-fold and used as template either for a 12.5 uL PCR reaction with Q5 Polymerase (NEB), or for a 10 uL qPCR reaction with the SsoAdvanced Universal SYBR Green 2X Supermix (BioRad). Primer pairs were designed as described previously (5). Briefly, each reaction includes one primer specific to the genome, and one primer specific to the mini-transposon, as shown in **Fig. 2b**.

For PCR, products were resolved by electrophoresis on 1.5% agarose gels stained with SYBR Safe (Thermo Scientific), and purified PCR products were confirmed by Sanger sequencing (Genewiz). For qPCR, each sample was analyzed using three primer pairs in three reactions: two as shown in **Fig. 2b** probing for the T-RL and T-LR orientations, respectively, and a third pair amplifying a reference genomic region. Integration efficiency (%) for each orientation is defined as 100 × (2^ΔCq), where ΔCq is the Cq(genomic reference pair) – Cq(T-RL pair OR T-LR pair); the total integration efficiency is the sum of both orientation efficiencies. This qPCR approach has been previously benchmarked using lysate samples simulating known integration efficiencies and orientation biases (5).

### SMRT-sequencing

Colonies were scraped from LB-agar plates after incubation and resuspended in LB media, and genomic DNA (gDNA) was extracted using the Wizard Genomic DNA Purification kit (Promega). Multiplexed, whole-genome SMRTbell libraries were prepared as recommended by the manufacturer (Pacific Biosciences). Briefly, 1 μg of high molecular weight gDNA from each sample (n=20–22 per pool) was sheared using a g-TUBE to ~15 kb (Covaris). Sheared gDNA samples were then used as input for SMRTbell preparation using the Express Template Preparation Kit 2.0, where each sample was treated with a DNA Damage Repair and End Repair/A-tail mix, in order to repair nicked DNA and create A-tailed ends. Barcoded overhang SMRTbell adapters were ligated onto each sample to complete SMRTbell library construction, and these libraries were then pooled equimolarly, with a final multiplex of 20–22 samples per pool. The pooled libraries were then cleaned up with 0.45X AMPure PB beads (Pacific Biosciences) and subjected to size-selection on Blue Pippin (SAGE Science) in order to remove DNA fragments <7 kb. The completed 20–22-plex pools were annealed to sequencing primer V4 and bound to sequencing polymerase 2.0, and were sequenced using one SMRTcell 8M on the Sequel 2 system, with a 30-hour movie.

After data collection, the raw sequencing reads were demultiplexed according to their corresponding barcodes using PacBio’s *Lima* tool (version 1.11.0). Circular consensus sequencing algorithm (CCS version 4.2.0) was used to perform intramolecular error correction on demultiplexed subreads with at least 3 polymerase passes, to generate highly accurate (>Q20) CCS reads. Each sample yielded between 5.9 and 12.6 Gb of total data (median = 9.9 Gb); CCS generation yielded between 28.2k and 81.6k CCS reads (median = 55.0k reads), with average CCS read lengths between 10.8 kb and 12.4 kb (median = 11.7 kb). For each sample, median CCS read quality ranged from Q32 to Q36. Full sequencing statistics are provided in **Supplementary Table 1**.

### SMRT-seq data analysis

Analysis of CCS reads from SMRT-seq was performed using a custom Python script. For each sample, BLASTn was performed on the CCS read sequences to search for the mini-transposon (mini-Tn) sequence. BLASTn hits that were not within 5-bp of the length of the mini-Tn, or that had E-values greater than 0.000001, were removed. Reads without any valid hits for the mini-Tn sequence were removed from further analyses; if multiple mini-Tn hits were identified in a read, the hits were analyzed separately in order of 5’-to-3’ position within the read.

For each mini-Tn hit within a read, 50-bp sequences flanking the mini-Tn were then extracted; flanking sequences shorter than 50 bp (i.e. mini-Tn hits positioned near either edge of the read) were discarded. This resulted in a list of 50-bp sequences flanking the mini-Tn for each read, with a pair of sequences corresponding to each mini-Tn hit. These flanking sequences (‘flanks’) were then classified as either donor-plasmid-mapping (‘plasmid flanks’) or genome-mapping (‘genomic flanks’), as follows: for each 50-bp flank, bowtie2 was used to align the flank to both the full donor plasmid sequence and the full target genome. The flank sequence was classified as genomic or plasmid based on which alignment had a lower Hamming distance, with a maximum allowed Hamming distance of 2. Flanks with Hamming distances greater than 2 from any genome or plasmid sequence were not classified, and the corresponding read was removed from further analyses. This process results in a list of classified flanks for each read, which was then used to classify the entire read. Reads containing only plasmid flanks were classified as plasmid reads; reads containing only genomic flanks were classified as simple insertion transposition products; and reads containing a mixture of plasmid and genomic flanks were classified as cointegrate transposition products.

The distance between multiple mini-Tn hits from reads categorized as cointegrates were manually inspected and confirmed to be the exact predicted length of the donor plasmid backbone. For all reads containing multiple mini-Tns (including ones not categorized as cointegrates or simple insertions), the same distance analysis was performed to look for mini-Tn hits within several bp from each other, which would be characteristic of tandem insertions; only one such read was found across all samples. We note that there were a number of observed reads that contained multiple consecutive plasmid flanks, suggesting the presence of concatemer plasmids and resulting concatemer cointegrates (23). These reads were classified using the same rules as above.

For each read that fell into either the simple insertion or cointegrate categories, the genomic coordinate where the genomic flank sequence mapped was recorded. For reads with more than one genomic-mapping flank, the first flank in the read (from 5’-to-3’) was used to determine the genomic insertion location. Extracted genomic coordinates were used to generate genome-wide histograms of integration locations; for visualization purposes, these locations were grouped into 912 5-kb bins. On-target reads were defined as reads with genomic insertion locations within a 100-bp window, centered at the site X-bp downstream of the 3’ end of the target site complementary to the guide RNA (5), with X=49 for the *V. cholerae* system and X=40 for the *S. hofmannii* system.

Alignments of exemplary CCS reads showing either simple insertions or cointegrates were performed using Geneious Prime 2020.2 at medium sensitivity with no fine-tuning. Simple insertion reads were aligned to a synthetic reference genome with a simple insertion product added at the expected target site; cointegrate reads were aligned to a synthetic reference genome with a cointegrate product added at the expected target site. Alignments were visualized using IGV 2.8.2, with indels < 10 bp not shown.

## Acknowledgements

We thank N. Jaber and S. E. Klompe for laboratory support, L.F. Landweber for qPCR instrument access, and I. Oussenko, N. Francoeur, and the Genomics Technology Laboratory at the Icahn School of Medicine at Mount Sinai for SMRT sequencing.

## Author Contributions

S.H.S. and P.L.H.V. designed the project and experiments, with input from M.S. P.L.H.V. performed all *E. coli* experiments; M.S. provided input and guidance on SMRT sequencing and analysis. C.A. developed custom algorithms to analyze sequencing data, with input from P.L.H.V. P.L.H.V., S.H.S., and all other authors discussed the data and wrote the manuscript.

## Availability of Data and Materials

Sequencing datasets are available at Sequence Read Archive NCBI Sequence Read (Accession-). Custom Python algorithms used for analysis are available at https://github.com/sternberglab/Vo_etal_2021. Datasets generated and analyzed during the current study are available from the corresponding author upon reasonable request.

## SUPPLEMENTARY FIGURES

**Supplementary Figure 1.**
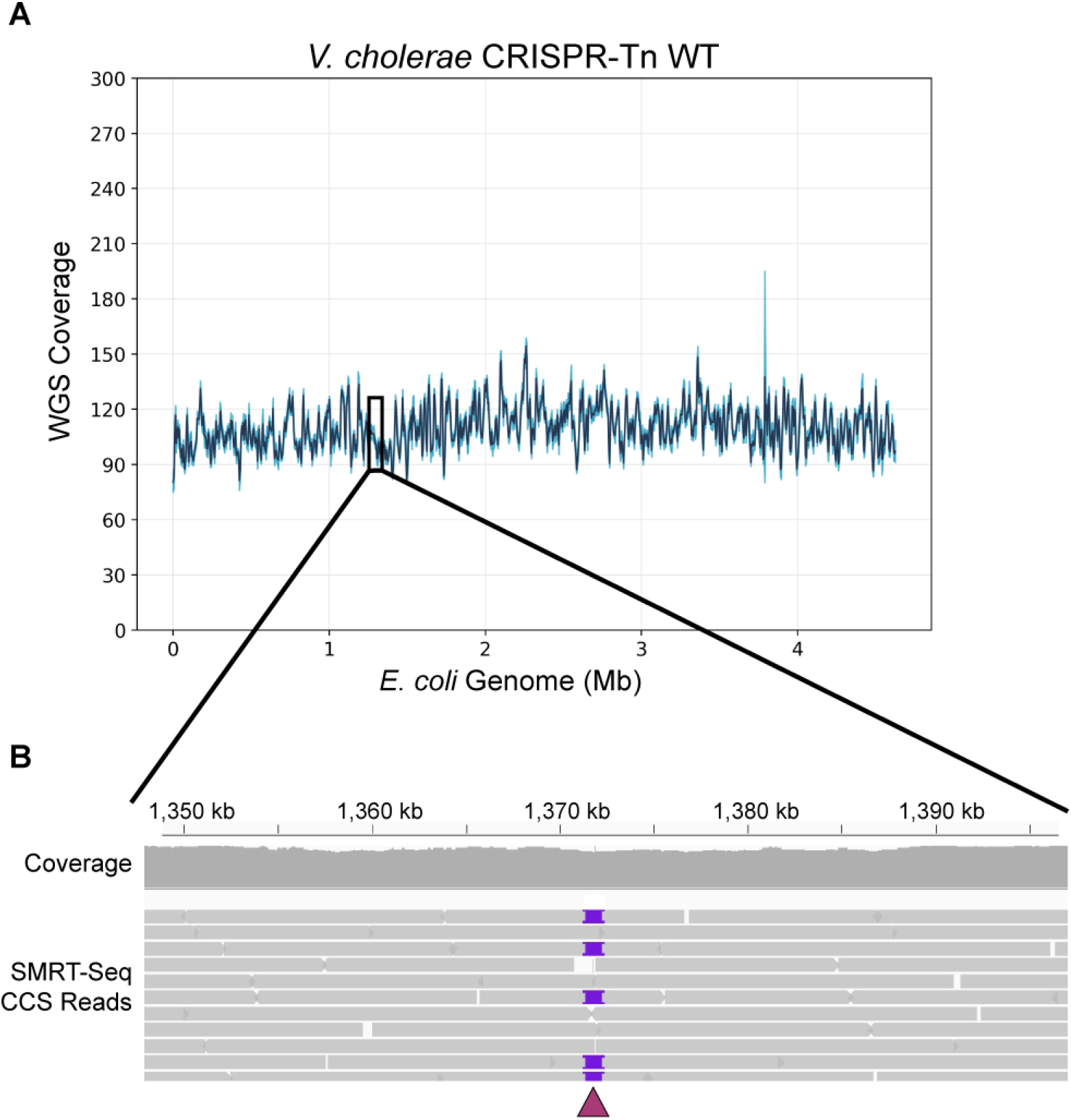
SMRT-seq approach to profile genome-wide transposition products. **a,** Representative read coverage from whole-genome SMRT-sequencing of *E. coli* strain BW25113, transformed with vectors encoding the WT *V, cholerae* CRISPR-transposon system. Read coverage was 90-120X coverage across the full-length genome. **b,** Representative CCS reads from whole-genome SMRT-seq data in **a**, aligned to the parental strain reference genome at the target site (maroon triangle). Purple blocks indicate large insertions corresponding to genomic RNA-guided DNA integration events.

**Supplementary Figure 2.**
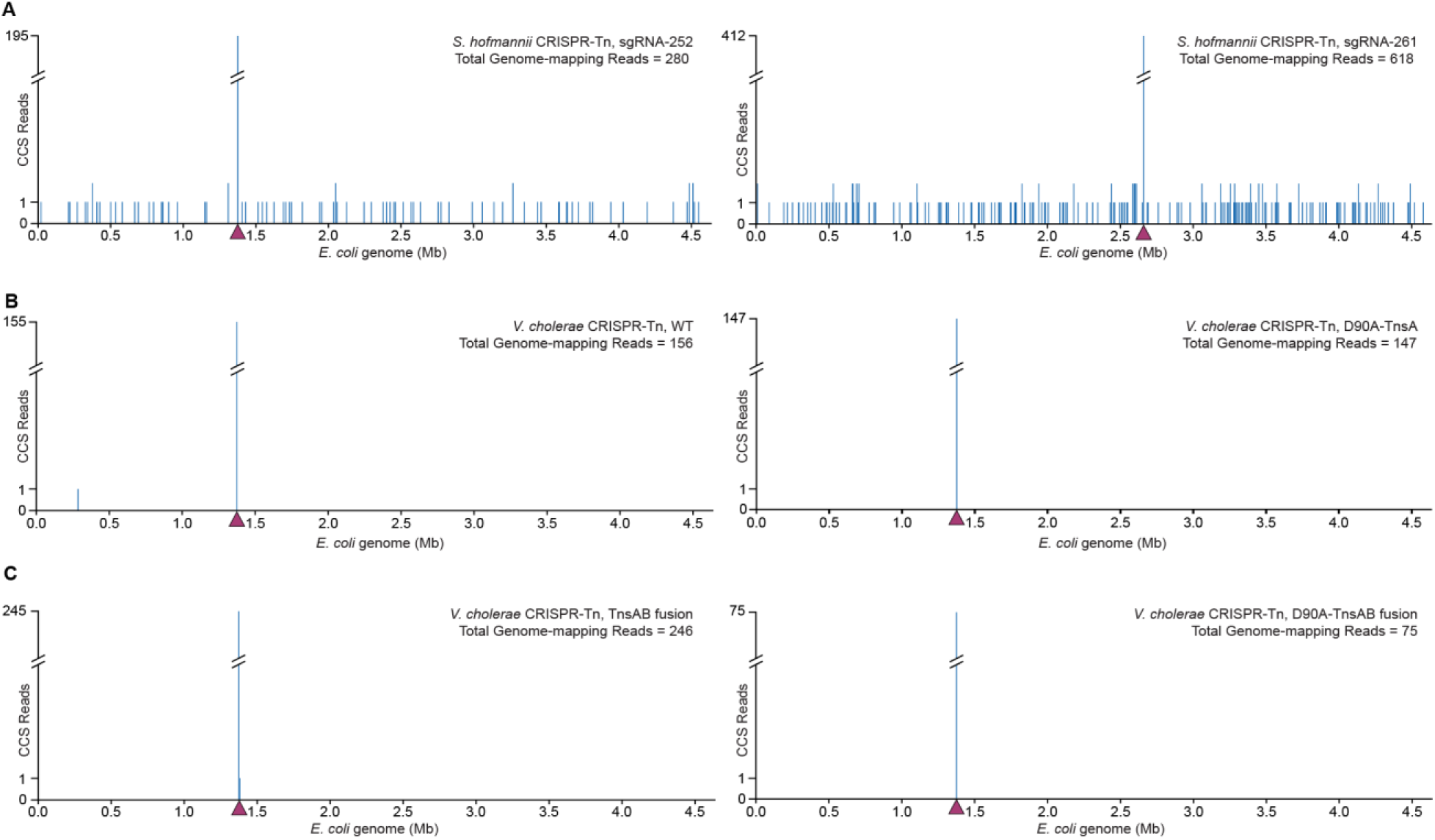
Genome-wide analyses of integration events extracted from SMRT-seq data. **a,** Integration events were determined from SMRT-seq CCS reads containing transposons flanked by genome-mapping sequences, for both simple insertion and cointegrate transposition products. Data are shown for the *S. hofmannii* CRISPR-Tn using either sgRNA-252 (left) or sgRNA-261 (right); target sites are denoted by maroon triangles. Note the scaling on the y-axis. **b,** Data for the *V. cholerae* CRISPR-Tn encoding WT (left) or D90A-TnsA (right), shown as in **a**. **c,** Data for the *V. cholerae* CRISPR-Tn encoding WT (left) or D90A-TnsAB fusion (right), shown as in **a**.

